# The functional organization of chromosome territories in single nuclei during zygotic genome activation

**DOI:** 10.1101/2025.04.06.647428

**Authors:** Akshada Shankar Ganesh, Taylor M. Orban, Romir Raj, Peter I. Fatzinger, Anna Johnson, Sean M. Riccard, Akhmed Zhanaidarov, Mayu Inaba, Jelena Erceg

**Affiliations:** Department of Molecular and Cell Biology, University of Connecticut, Storrs, CT 06269, USA; Institute for Systems Genomics, University of Connecticut, Storrs, CT 06269, USA; Department of Cell Biology, University of Connecticut Health Center, Farmington, CT 06032, USA; Department of Genetics and Genome Sciences, University of Connecticut Health Center, Farmington, CT 06030, USA

**Keywords:** chromosome territories (CTs), transcription, homolog pairing, zygotic genome activation (ZGA), haploid, RNA polymerase II

## Abstract

Chromosome territories (CTs) are intricately organized and regulated within the nucleus. Despite remarkable advances in our understanding of genome packaging and gene expression, the interplay among CTs, pairing of parental homologous chromosomes, and genome function during development remains elusive. Here, we employ an Oligopaints-based high-resolution imaging approach to examine variable CT organization in single nuclei during the developmental process of zygotic genome activation. We reveal large-scale chromosome changes with extensive homolog pairing at the whole-chromosome level that decreases locally due to spatial variability in chromosome conformations. In the absence of one homolog copy, the dynamics of CT compaction and RNA polymerase II recruitment are supported by transcriptional changes in haploid embryos. Finally, global inhibition of transcription results in decreased CT opening and no significant impact on CT pairing levels. These findings enhance our understanding of parental genome folding and regulation, which may inform strategies for chromosome-based diseases.

## Introduction

Individual chromosomes largely occupy discrete areas within the nucleus leading to the formation of independent chromosome territories^1-4^. While such intricate chromosome packaging has been observed by population-based approaches, such as derivatives of chromosome conformation capture^5-11^, and single-cell imaging^2,3,12-18^, studies investigating cell-type variability of CTs during development remain limited. Proper chromosome integrity is essential as aberrant chromosomal alterations may be related to dysfunctional chromatin and lead to developmental disorders^19,20^. Copy number alterations such as chromosomal gains and losses are pervasive in cancer^21^. Although most interactions are intra-chromosomal within individual chromosomes, the spatial proximity between different chromosomes in *trans* can be associated with intermingling of neighboring CTs and translocations with significant bearing on gene regulation^1,13,16,22-35^. Interestingly, this proximity may involve pairing of the homologous maternal and paternal chromosomes. These *trans*-homolog interactions are not only restricted to meiosis, they may also occur in somatic cells and increase as development progresses in *Drosophila*^5,36-46^. Such homolog pairing exhibits a high degree of structural organization which may bear functional implications to chromatin activity and transcription at a global scale^5,46-49^. Pairing is also observed in mammalian systems related to imprinting, DNA repair, V(D)J recombination, cell fate, and X-chromosome inactivation^37,39,45^. Nevertheless, the broader impact of this *trans*-homolog organization in early development on CT compaction and regulation remains elusive.

During the initial developmental stages in diploid organisms, embryos rely on maternally-contributed products. As the embryonic genome awakens, zygotic transcription begins in a process called zygotic genome activation (ZGA)^50-52^. This transcription occurs in two different stages, the minor and major wave of ZGA. For instance, the onset of the minor wave of ZGA in *Drosophila* is marked by the production of the early transcripts during nuclear cycle 8, and the major wave of ZGA is characterized by large-scale transcriptional activation at nuclear cycle 14^50-52^. Furthermore, during ZGA additional events occur such as global RNA polymerase II (RNA Pol II) recruitment^6,53-56^, widespread increase in chromatin accessibility^57-65^, and structural genome remodeling^6,8,66,67^. However, the interplay of transcriptional activation with homolog pairing and chromosome packaging in single nuclei of developing embryos is still unclear.

Here, we asked how CTs may be impacted by homolog pairing and transcription on a global scale during ZGA. To achieve that, we turned to *Drosophila* as a powerful model system with only 4 homologs, abundant embryos, and prominent somatic homolog pairing^37,39,45^. We employed a customized Oligopaints-based imaging approach^68,69^ to visualize spatial heterogeneity of CTs across the entire genome in developing embryos during the onset of ZGA. We reveal changes in genome folding at the whole-chromosome and chromosome-arm scale. Moreover, at the whole-chromosome scale, parental homologs show extensive pairing in single nuclei, while pairing at the chromosome-arm level becomes less precise, suggesting spatial variability in chromosome conformations. When comparing the absence of one homolog copy in haploid embryos to diploid counter parts, we find that variations in chromosome compaction and RNA Pol II recruitment may be supported by changes to transcriptional output. Conversely, transcription inhibition in developing embryos results in decreased CT opening and does not significantly impact levels of CT pairing. Together, these findings provide an enhanced framework for our understanding of global parental genome folding and regulation during early embryogenesis.

## Results

### Large-scale chromosome changes during ZGA in single nuclei of developing embryos

To examine the dynamics of CTs, we took advantage of a crucial period when homologs pair and the genome awakens during embryogenesis^41-43,50-52^. In particular, we focused on the onset of the minor and major waves of ZGA, corresponding to nuclear cycles 8 and 14, respectively (Figure 1A). In the minor wave, only a small portion of genes are expressed, while global gene activation leads to the major wave of ZGA^50-52^. To characterize CTs, we utilized the Oligopaints approach^68,69^ to target chromosomes X, 2, 3, and 4, excluding repetitive regions. We designed 299,701 specialized oligos with probe density ≥ 2.30 probes/kb using the OligoMiner setting (Figure 1B; Table S1; Methods). We successfully applied these customized probes to visualize CTs of 2, 3, and 4 in developing *Drosophila* embryos (Figures 1C and 1D). Formation of such independent CTs is supported by *Drosophila* embryonic Hi-C^5,6,8-11^ and imaging data of various cell types^2,3,12,15,16^. In addition, to distinguish chromosome X in female versus male embryos, we utilized a satellite repeat probe (AATAT)_n_^70-72^. Since this repeat probe primarily labels highly repetitive chromosomes Y and 4, we combined it with our chromosome 4 Oligopaint probe to discern the presence of Y (Figures S1A and S1B).

**Figure 1.**
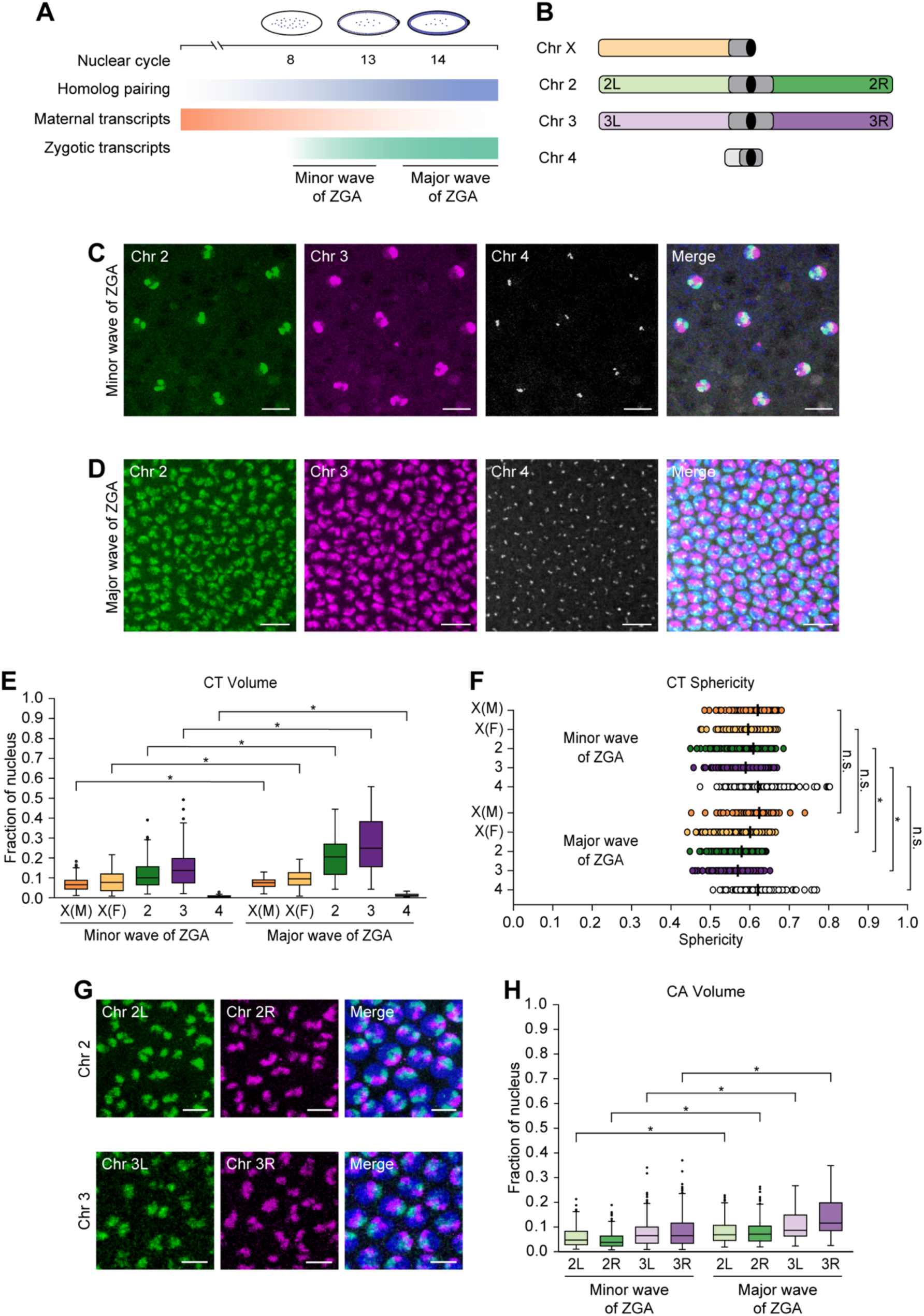
Changes in CTs and CAs during the minor and major waves of ZGA. (A) Schematic representation of events during the onset of minor (nuclear cycle 8) and major (nuclear cycle 14) waves of ZGA. Maternal transcripts (orange) decrease, while zygotic transcripts (green) and somatic homolog pairing (blue) progressively increase. (B) Oligopaint probes target *Drosophila* chromosomes X, 2, 3, and 4 along with the arms of major chromosomes labelled. Centromere, black; heterochromatin, dark gray. (C and D) Chromosomes 2 (green), 3 (magenta), and 4 (gray) of *Drosophila* embryos during the minor (C) and major (D) waves of ZGA. Total DNA by Hoechst stain (blue). Bar = 10 μm. (E) CT volume from the minor to major waves of ZGA. Each CT volume is normalized to the respective nuclear volume. X(M), chromosome X in males; X(F), chromosome X in females; at least three replicates; n ≥ 300 nuclei; *p ≤ 1.27×10^-6^, Mann-Whitney two-sided *U* test. (F) CT sphericity between the minor and major waves of ZGA. X(M), chromosome X in males; X(F), chromosome X in females; median, solid line; at least three replicates; n ≥ 300 nuclei; *p ≤ 7.33×10^-13^, n.s., not significant, Mann-Whitney two-sided *U* test. (G) Representative images display the left (green) and right (magenta) arms of chromosomes 2 (top) and 3 (bottom) in the major wave of ZGA. Total DNA by Hoechst stain (blue). Bar = 5 μm. (H) CA volume from the minor to major waves of ZGA. Each CA volume is normalized to the respective nuclear volume. At least three replicates; n ≥ 300 nuclei; *p ≤ 1.94×10^-10^, Mann-Whitney two-sided *U* test.

Following microscopy using the Oligopaint probes, we generated a computational pipeline to quantify chromosome dimensions and morphology in individual segmented nuclei (Methods). Previous studies using genome-wide assays such as MNase-seq, DNaseI sensitivity, and ATAC-seq observed increased chromatin accessibility during ZGA^57,58,60,63,64^. Similarly, using our image analysis pipeline at single-nucleus resolution, we observed a significant increase in the normalized volumes of chromosomes X, 2, 3, and 4 during the major wave compared to minor wave of ZGA (p ≤ 1.27×10^-6^) (Figure 1E; Table S1). In addition, we quantified the sphericity of CTs, a measure of how closely their shape approximates to an ideal sphere (Figure S1C). As CT volumes increase, we noticed that the sphericity of all chromosomes follow a similar decreasing trend, with significance only for autosomes 2 and 3 (p ≤ 7.33×10^-13^) (Figure 1F; Table S1). Since chromatin folding is related to epigenetic states where large volumes with less sphericity are associated with active chromatin^73,74^, these changes in CT compaction may reflect chromatin opening.

To explore this further, we utilized our Oligopaint probes to differentialy label the arms of major chromosomes 2 and 3 to determine if these trends are also consistent for individual chromosome arms (CAs) (Figures 1B and 1G). Comparable to CTs, as the zygotic genome is activated, we observed a significant increase in the normalized volumes of arms 2L, 2R, 3L, and 3R (p ≤ 1.94×10^-10^) (Figure 1H; Table S1). Although not significant (p ≤ 0.542), we observed a similar trend with the sphericity of CAs, suggesting changes in compaction based on the opening of the chromatin (Figure S1D; Table S1). In summary, our data reveals single-nucleus changes in large-scale genome packaging at the whole-chromosome and chromosome-arm level as developing embryos progress from the minor to major wave of ZGA.

### Whole-chromosome scale homologs are extensively paired in single nuclei

Since homologous chromosomes pair at many individual loci throughout embryogenesis as revealed by imaging and Hi-C^5,38,41-44^, we systematically investigated single-nucleus homolog pairing in the global context of CTs and CAs during ZGA (Figure 2A). Excitingly, using high-resolution microscopy, at the scale of whole chromosomes, X in females, 2, and 3, homologs are very highly paired appearing often as a single signal (70.6% ± 6.5–83.0% ± 1.0) (Figure 2B; Table S2). Though homologous chromosomes may spatially come together in many conformations, not all of them are amenable to pairing of CAs, leading to decreased arms pairing (36.2% ± 6.8–43.0% ± 0.5) (Figures 2C and 2D). In contrast to those major chromosomes with over 20 megabases (Mb) per arm, the pairing of the smaller chromosome 4 of 1.35 Mb is lower (26.6% ± 7.6–41.0% ± 4.2) (Figure 2B; Table S2). Similarly, previously studied individual euchromatic loci ranging with hundreds of kilobases (kb) have an even lower range of pairing (∼1–30%)^5,38,41-44^. Together, these observations suggest that, at the whole-chromosome scale, homologs may be extensively paired in single nuclei; however, that pairing is less precise and not well-aligned locally.

**Figure 2.**
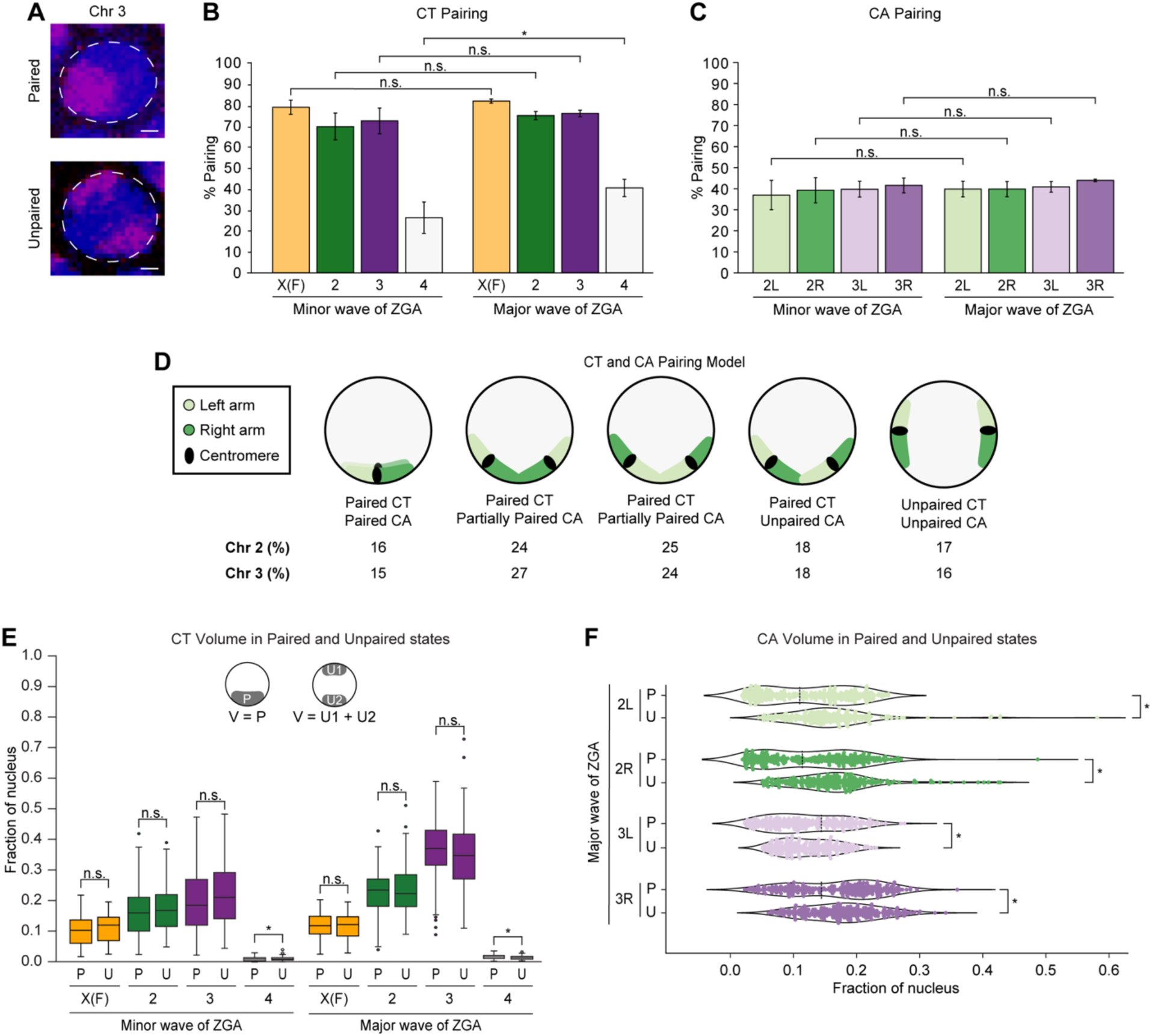
Extensive homolog pairing of CTs and CAs during ZGA. (A) Paired (top) and unpaired (bottom) chromosome 3 (magenta) during major wave of ZGA. Total DNA by Hoechst stain (blue). Bar = 1 μm. Percentage of nuclei showing (B) CT pairing and (C) CA pairing from the minor to major waves of ZGA. X(F), chromosome X in females; error bars, standard deviation; at least three replicates; n ≥ 300 nuclei; *p = 6.26×10^-3^, n.s., not significant, Fisher’s two-tailed exact test. (D) Schematic representation illustrating the percentage of occurrence in major wave of ZGA for the different conformations of left (light green) and right (dark green) arm, indicating higher CT than CA pairing. The angles between CAs may vary. (E) Normalized CT volume between paired homologs to the combined volume of two unpaired homologs from the minor and major waves of ZGA. X(F), chromosome X in females; P, paired; U, unpaired; U1, unpaired homolog 1; U2, unpaired homolog 2; V, volume; at least three replicates; n ≥ 88 nuclei; *p ≤ 8.89×10^-7^, n.s., not significant, Mann-Whitney two-sided *U* test. (F) Normalized CA volume differences in arms of paired homologs to the combined volume of two unpaired homologs during the major wave of ZGA. P, paired; U, unpaired; local minima, dashed line; at least three replicates; n ≥ 242 nuclei; *p ≤ 3.06×10^-5^, n.s., not significant, Levene’s test.

We next investigated if the pairing alters the compaction of chromosomes as the genome awakens. We observed that paired homologs occupy a significantly higher fraction of the nucleus compared to individual unpaired homologs (p ≤ 1.26×10^-9^) (Figure S2A; Table S2). Likewise, arms of paired homologs are more open than arms of individual unpaired homologs (p ≤ 2.67×10^-4^) (Figure S2B; Table S2). While we noticed slight differences in the combined volume of two unpaired homologs to the paired homologs of chromosomes X in females, 2, and 3, these differences are largely negligible (p ≤ 0.596), suggesting no significant changes in total chromatin volume based on the pairing of homologs (Figure 2E; Table S2). Interestingly, in contrast to the minor wave of ZGA (Figure S2C), we discerned two distinct populations within arms of paired homologs in the major wave of ZGA: one characterized by tight pairing and another by loose pairing (Figure 2F) suggesting associations with genome activity and silencing, respectively^5,47^. Furthermore, during the minor wave of ZGA, the sphericities of chromosomes X in females, 2, and 3, as well as their corresponding arms are variable. As the genome awakens during the major wave, unpaired homologs have significantly lower sphericity than paired homologs (p ≤ 3.58×10^-2^) (Figures S2D and S2E; Table S2). Altogether, these observations suggest that while extensive homolog pairing does not lead to large-scale volume changes, it may significantly influence the shape of the chromosomes.

### Variations in CT compaction and RNA Pol II recruitment are supported by changes in transcriptional levels in haploid embryos

To further understand the impact of *trans* interactions between homologs in the diploid genome, we leveraged homozygous *maternal haploid* (*mh*) females to produce haploid embryos, which eliminates *trans*-homolog interactions entirely. In addition, it reduces the copy number of chromosomes from 2n to n. The *mh* haploid embryos can develop until late embryogenesis and never hatch. These embryos also have smaller nuclei than diploid embryos, and undergo an additional round of nuclear division during the blastoderm stage (Figure 3A)^75-81^. Consistent with previous studies^75,78,79^, the absence of one homolog copy in haploid embryos led to significantly lower nuclear volume compared to diploid embryos (p ≤ 2.09×10^-10^) (Figure 3B; Table S3). During nuclear cycle 14 in haploid embryos, the CT volumes normalized per nuclear volume are significantly higher than their individual unpaired counterparts in diploid embryos, but a sharp decrease in the volumes is observed at nuclear cycle 15 (p ≤ 1.81×10^-2^) (Figure 3C; Table S3). This trend may be attributed to the transcriptional output of haploid embryos, where time-dependent genes show hyperactivity with increased RNA levels compared to diploid embryos during nuclear cycle 14^78^. However, by late cycle 14 this hyperactivity disappears. In haploids at nuclear cycle 15, a small number of genes sensitive to the ratio of nuclear content to cytoplasmic volume (N/C ratio) are activated, but this transcription may not be sufficient for global decompaction. Additionally, we observed that the corresponding CT sphericity in diploid versus haploid embryos follows a similar pattern as the CT volume, further supporting observations that pairing influences the shape of the chromosomes (p ≤ 1.87×10^-7^) (Figure S3A; Table S3).

**Figure 3.**
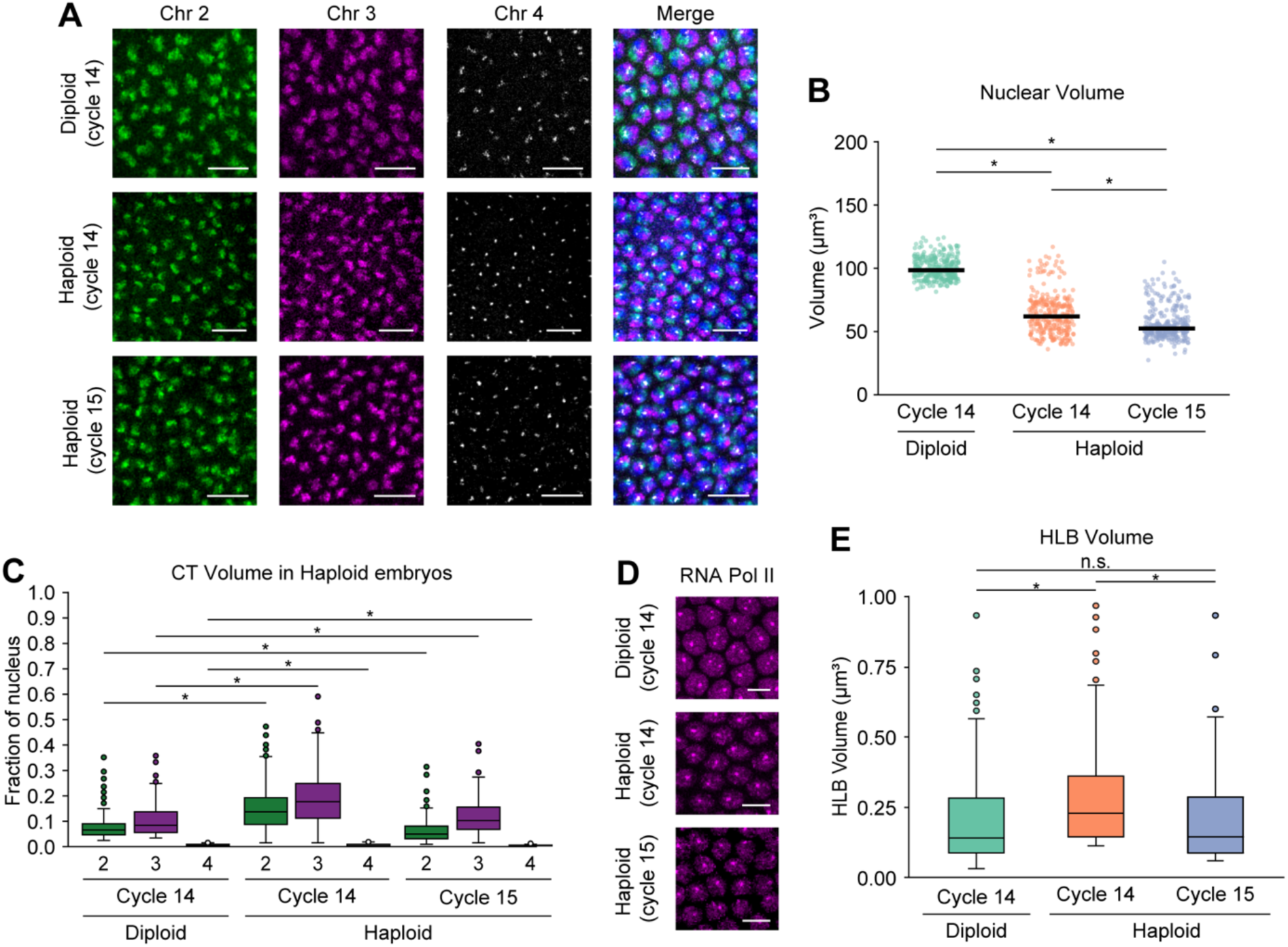
CT dynamics in haploid embryos during ZGA. (A) Chromosomes 2, 3, and 4 in diploid (cycle 14, top) and haploid embryos (cycle 14, middle; cycle 15, bottom) during major wave of ZGA. Total DNA by Hoechst stain (blue). Bar = 10 μm. (B) Nuclear volume between diploid (cycle 14) and haploid embryos (cycles 14 and 15). Median, solid line; at least three replicates; n ≥ 300 nuclei; *p ≤ 2.09×10^-10^, Mann-Whitney two-sided *U* test. (C) CT volume differences for chromosomes 2, 3, and 4 in diploid and haploid embryos. In diploid embryos, only individual unpaired homolog volumes were used. Each CT volume is normalized to the respective nuclear volume. At least three replicates; n ≥ 300 nuclei; *p ≤ 1.81×10^-2^, Mann-Whitney two-sided *U* test. (D) RNA Pol II in diploid (cycle 14, top) and haploid embryos (cycle 14, middle; cycle 15, bottom) during major wave of ZGA. Bar = 5 μm. (E) HLB volume between diploid and haploid embryos. In diploid embryos, only individual unpaired homolog volumes were used. Median, solid line; at least three replicates; n ≥ 300 nuclei; *p ≤ 3.20×10^-15^, n.s., not significant, Mann-Whitney two-sided *U* test.

We next examined how the absence of one homolog copy affects transcription in single nuclei of haploid embryos. To this end, we used high-resolution microscopy to visualize the RPB1 subunit of RNA Pol II, targeting all forms of RNA Pol II in haploid and diploid embryos (Figure 3D). In diploid embryos, we observed that RNA Pol II is recruited globally across nuclei; however, two distinct foci are larger than any other RNA Pol II foci (Figure 3D). These correspond to histone locus bodies (HLBs), which play a role in biosynthesis and processing of histone mRNAs^55,82,83^. As expected, only one focus is found in haploid embryos, in agreement with copy number difference (Figure 3D). As higher RNA Poll II accumulation in HLBs may lead to higher expression of core histone genes^55^, we inspected whether HLB volume follows the same trend as CT volume. We found that HLB volume increases from diploid to haploid embryos in cycle 14, then in haploid it decreases from cycle 14 to cycle 15 (p ≤ 3.20×10^-15^) (Figure 3E; Table S3). Similar to CT volume of haploid embryos, this variation may also be supported by higher transcriptional output in haploid compared to diploid embryos^78^. Excluding the signal at the HLBs, the RNA Pol II intensity significantly decreases from diploid to haploid embryos (p ≤ 3.33×10^-18^) (Figure S3B; Table S3) due to the presence of twice the DNA template content in diploid than haploid embryos. Together, with the absence of one homolog copy in haploid embryos, structural CT compaction and RNA Pol II recruitment relate to transcriptome changes.

### Transcription inhibition impacts CT compaction, but not pairing levels at the whole-chromosome scale

Transcription inhibition may impact domain boundary insulation and condensate formation^6,84-86^. Hence, we next utilized high-resolution microscopy to investigate if transcription may influence CT compaction and homolog pairing in single nuclei during *Drosophila* ZGA. To this end, we injected embryos with RNA Pol II inhibitors, alpha-amanitin and triptolide, and then collected them to capture the onset of the major wave of ZGA (Figure 4A). Nuclear volumes remain consistent between water-treated control embryos and those treated with RNA Pol II inhibitors (Figure 4B; Table S4). With transcription inhibition, the normalized volumes of chromosomes 2, 3, and 4 significantly decreased compared to the control (p ≤ 6.11×10^-3^) (Figure 4C; Table S4). Given the relationship between epigenetic states and chromatin folding^73,74^, our observations suggest that transcription inhibition may result in decreased chromatin opening. This is also corroborated with our observation of CT opening from the minor to major wave of ZGA (Figure 1E) as gene expression increases from a small fraction of genes to widespread gene activation.

**Figure 4.**
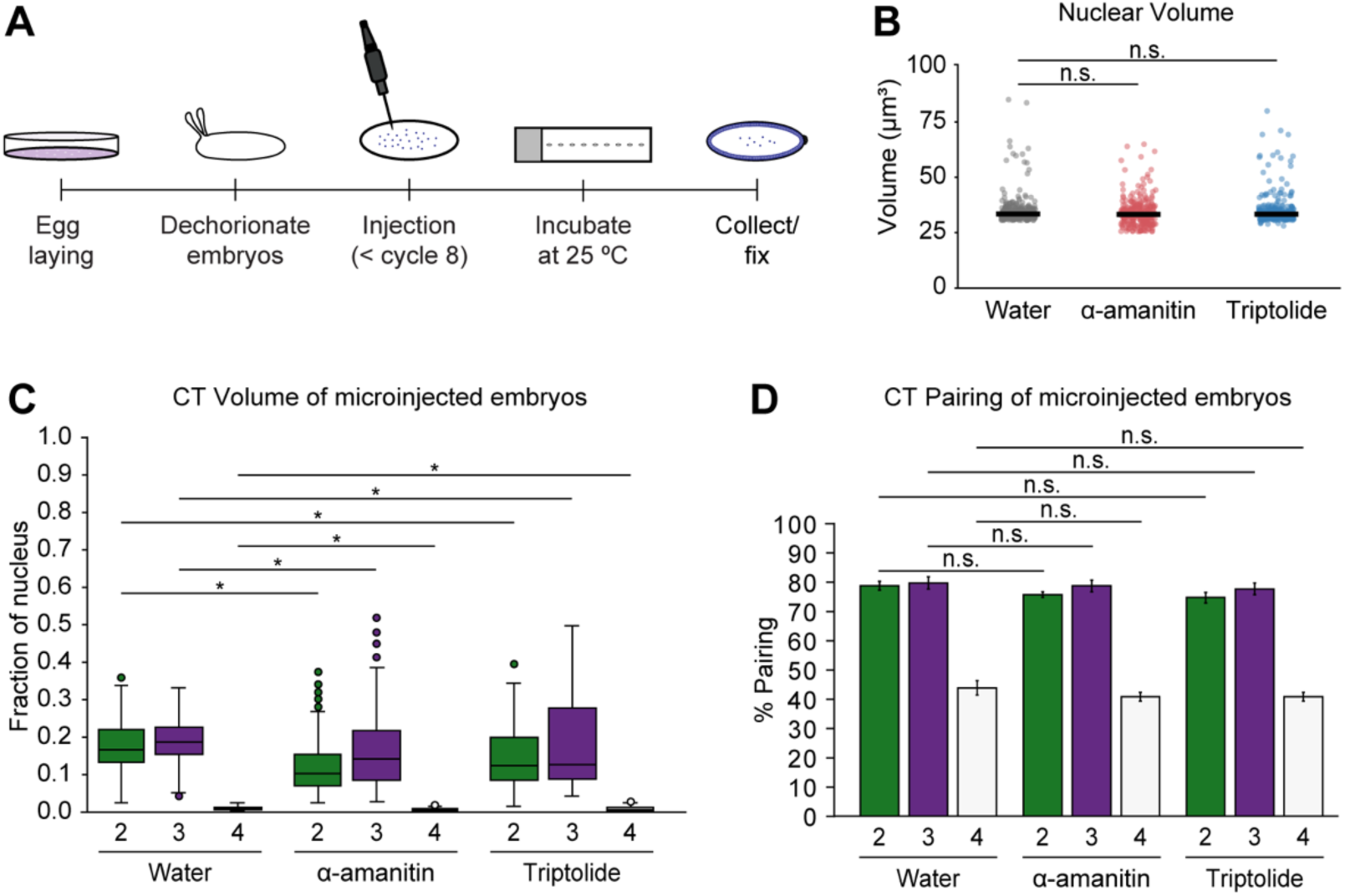
CTs and homolog pairing in transcription inhibited embryos. (A) Diagram illustrating embryonic microinjections. (B) Nuclear volume in control, alpha-amanitin, and triptolide-treated embryos. Median, solid line; at least three replicates; n ≥ 300 nuclei; n.s., not significant, Mann-Whitney two-sided *U* test. (C) Normalized CT volume for chromosomes 2, 3, and 4 in water (control), alpha-amanitin, and triptolide-treated embryos. At least three replicates; n ≥ 300 nuclei; *p ≤ 6.11×10^-3^, Mann-Whitney two-sided *U* test. (D) CT pairing of transcription inhibited and control embryos. Error bars, standard deviation; at least three replicates; n ≥ 300 nuclei; n.s., not significant, Fisher’s two-tailed exact test.

Next, we examined whether transcription inhibition affects homolog pairing. Our results reveal that, at the CT scale, homologs of major chromosomes remain highly paired (75.0% ± 1.7– 80.0% ± 2.0) and show no significant differences in pairing levels between water-treated and transcription inhibited embryos (Figure 4D; Table S4). Similarly, with the increase of transcription from the minor to major wave of ZGA, pairing levels of major chromosomes also exhibit no significant change (Figure 2B). Taken together, these findings support the connection between transcription and structural CT organization as well as suggest that transcription may not impact pairing levels at the global CT scale.

## Discussion

Our study reveals the dynamics of CTs, including intra- and inter-chromosomal interactions, and their impact on genome function using customized Oligopaint probes at the single-nucleus resolution. During the onset of ZGA, we uncover large-scale genome folding changes at both the whole-chromosome and chromosome-arm levels in *Drosophila* embryos. Our findings using high-resolution microscopy suggest that homologs may pair extensively at the whole-chromosome scale, while variable chromosome conformations may lead to less precise pairing locally. By eliminating homolog pairing in haploid embryos, we assessed the impact of *trans*-homolog organization on genome structure and function. We found that the dynamics of CT folding and recruitment of RNA Pol II associate with transcriptional output of haploid embryos with half the DNA template content. Meanwhile, perturbing transcription in diploid embryos using RNA Pol II inhibitors resulted in decreased CT opening, further corroborating the connection between transcription and CT organization. However, transcription inhibition did not significantly impact CT pairing levels.

Consistent with previous *Drosophila* embryonic Hi-C and imaging studies across multiple cell types^2,3,5,6,8-12,15,16^, we demonstrate for the first time the formation of independent CTs by individual chromosomes and chromosome arms in single nuclei of embryos during ZGA. Furthermore, our study reveals extensive somatic homolog pairing at the whole-chromosome scale, which can bear functional implications on zygotic gene expression through transvection^37,39,40,45^. Several models have been proposed to understand the underlying structure of somatic homolog pairing, including well-aligned chromosomes (railroad track), loose association of homologous regions (laissez-faire), button model, and highly disordered pairing^5,37,45,46,87^. Moreover, haplotype-resolved Hi-C studies found that homolog pairing has a highly structured organization with at least two forms of pairing, tight and loose^5,47^. Here, using high-resolution microscopy, we observed less precise pairing at the chromosome-arm level than at the whole-chromosome scale, where tight and loose pairing is associated with genome activity^5,47^. Notably, our CT and CA pairing model suggests multiple conformations and angles between chromosomes and chromosome arms. However, not all of these conformations are amenable to pairing, leading to reduced, less well-aligned local pairing. Together, these chromosome conformations indicate that intra- and inter-chromosomal organization is variable and dynamic across individual nuclei.

Although the genome structure is intricately organized, it remains elusive whether such 3D genome structure mirrors gene regulation or if genome architecture instructs gene expression^39,88-91^. Structural rearrangements can have drastic effects in modulating gene expression in disease and evolution^20,88,90^. However, some instances of chromosomal rearrangements and perturbation of factors implicated in genome organization do not result in widespread shifts in transcriptional output^20,88,90,92-95^. An additional layer of complexity to these contrasting observations is brought by inter-chromosomal interactions, which are implicated in translocations, transvection, acquisition of cellular identity, centromere and telomere clustering, and nuclear hub formation^37,39,40,45,96^. Specifically, finer details of the relationship between CT architecture, pairing, and genome regulation in single nuclei during early development remain elusive. In the absence of homolog pairing in haploid embryos, our findings suggest that pairing may be dispensable for the relationship of CT compaction and the recruitment of RNA Pol II, potentially due to changes in transcriptional outputs. With this copy number change of DNA template, RNA levels are altered^78^ as well as the associated levels of RNA Pol II recruitment and chromosome compaction. Previous studies suggest that cluster formation by RNA Pol II occupancy and domain boundary insulation may be affected by transcription inhibition^6,84-86,97^. Using high-resolution microscopy, our findings indicate that transcription inhibition leads to increased CT compaction and no significant effect on CT pairing, thereby, providing insight into the relationship between CTs, homolog pairing, and transcription.

The variable genome organization across cell populations during development may have functional implications in diseases^98,99^. Such variation at the whole-chromosome and chromosome-arm level may be associated with translocations and aneuploidy, which can lead to detrimental effects in cancer and developmental disorders^19-21^. Together, understanding genome-wide organizations of CTs and CAs as well as their association with transcription may benefit the strategies to combat chromosome-based diseases.

## Materials and Methods

### Collection and fixation of embryos

Hand-sorted, inbred virgin females from *Drosophila melanogaster* Genetic Reference Panel^100^ DGRP-057 (Bloomington stock number 29652) and males from DGRP-439 (Bloomington stock number 29658), that differ by many single nucleotide variants (SNVs), were crossed to obtain the F1 hybrid embryos. The genotype of these F1 hybrid embryos matches that of embryos used for haplotype-resolved Hi-C^5^ to facilitate comparison between imaging and Hi-C approaches. Nuclear cycles 8 and 14 embryos were collected following three pre-lays at 25 °C. Embryo fixation was performed as previously described^5,44,101^. Briefly, embryos were dechorionated with 50% bleach for 3 min and washed in 1x PBS with 0.1% Triton X-100. The embryos were shaken for 30 min in 500 µl of 4% paraformaldehyde, 0.5% Nonidet P-40, and 50 mM EGTA in 1x PBS, and 500 µl of heptane. The fix was replaced with methanol, followed by vigorous shaking for 1 min and three subsequent washes with 100% methanol. The embryos were then stored at −20 °C in methanol.

### Design and synthesis of FISH probes

The libraries for CTs and CAs were designed using the previously described Oligopaints approach with OligoMiner^68,69^. These libraries were purchased from Twist Bioscience and contain 299,701 specialized oligos with probe density ≥ 2.30 probes/kb (Table S1). Specific primers for these Oligopaint probes are provided in Table S1. As previously described^5,101^, forward primers were added a site for secondary oligo annealing, reverse primers were added a T7 promoter sequence, and secondary oligos contained both 5’ and 3’ conjugated fluorophores. An oligo probe for satellite repeat (AATAT)_n_^70-72^ was ordered from Integrated DNA Technologies (IDT) with this sequence and a fluorescent dye: /5Alex488N/AATATAATATAATATAATATAATATAATAT. Oligopaint probes were synthesized by T7 amplification with modifications from previous work^5,102^. The designed library was amplified using Kapa Taq enzyme (Kapa Biosystems, 5 U/µl), with the following PCR program: 95 °C, 5 min, (95 °C for 30 s, 58 °C for 30 s, 72 °C for 20 s) repeated 12 times, 72 °C 5 min, hold at 4 °C. The linear PCR products were purified using DNA Clean & Concentrator-5 (DCC-5) kit (Zymo Research). Linear PCR was followed by another bulkup PCR with 0.8 µM final concentration of the forward (with the site for secondary oligo annealing) and reverse (with T7 promoter sequence) primers, again purified using the same kit. Following purification, T7 RNA polymerase mix (HiScribe T7 High Yield RNA Synthesis Kit, NEB) and RNAse OUT (ThermoFisher Scientific) were added to the purified PCR product to produce excess RNA at 37 °C. Using the reverse transcriptase Maxima H Minus RT (ThermoFisher Scientific), RNA was reverse transcribed into DNA, and the RT enzyme was inactivated at 85 °C for 5 min. After the inactivation of the RT enzyme, all RNA in the solution was degraded using alkaline hydrolysis (0.5 M EDTA and 1 M NaOH in 1:1) at 95 °C for 10 min. Subsequently, the oligos were purified using the same clean-up kit with the Oligo binding buffer (Zymo Research).

### DNA FISH in whole embryos

DNA fluorescent in situ hybridization (FISH)^5,44,101^ was conducted in whole *Drosophila* embryos with the following modifications. Fixed embryos were rehydrated in succession from 100% methanol to 2x SSCT (0.3 M NaCl, 0.03 M sodium citrate, 0.1% Tween-20) at room temperature (RT), followed by two quick washes and a 10-minute wash in 2x SSCT. The embryos were incubated for 10 min in 2x SSCT/20% formamide and subsequently another 10 min in 2x SSCT/50% formamide. The primary hybridization buffer (2x SSCT, 10% dextran sulphate, 50% formamide, RNase A) along with Oligopaint probes were added to the embryos and incubated for 30 min at 80 °C, and then left overnight at 37 °C. For major chromosomes 2 and 3, 400 pmol of each arm probe was added to the hybridization buffer. For FISH targeting chromosome 4 and specific arms, 200 pmol of Oligopaint probes were used. To distinguish chromosome X in males and females, 200 pmol of the chromosome X Oligopaint probe and 200 pmol of a satellite repeat probe (AATAT)_n_ were added to the primary hybridization mix. Upon primary probe hybridization, the embryos were washed for 30 min in 2x SSCT/50% formamide at 37 °C. Following this, embryos were incubated for 30 min with 200 pmol of secondary probes containing fluorophores in 2x SSCT/50% formamide at 37 °C. The embryos were washed for 30 min in 2x SSCT/50% formamide at 37 °C, then for 10 min in 2x SSCT/20% formamide at RT, and quickly rinsed twice in 2X SSCT. Hoechst 33342 (1:1000, Invitrogen) was added to a third 2x SSCT rinse and incubated for 10 min at RT. Another wash of 10 min in 2x SSCT was followed by a quick rinse in 2x SSC. The embryos were then mounted in SlowFade Gold antifade reagent (Invitrogen) for imaging.

### Imaging data acquisition and analysis

Images with Z-stacks were acquired using a Leica SP8 confocal microscope with a 63x/1.4 oil-immersion objective lens at 1024×1024 resolution. Images were segmented and quantified for volume and sphericity using a custom pipeline on ZEISS arivis Pro Software, version 4.1.0. The analysis pipeline used the ‘Blob Finder’ feature for segmentation and the ‘Compartments’ feature to identify individual signal in respective nuclei. To determine the 3D distance between two FISH signals, each Z-stack was manually examined using the Fiji (ImageJ2, Version 2.16.0/1.54g) software, and the point tool was used to determine the x, y, and z coordinates. Homologs were defined as paired if the 3D distance between two signals was ≤0.8 µm or if only one FISH signal was present. RNA Pol II intensity was calculated using Measure feature. The nuclei that were counted for Figures 2C and 2D may vary.

### Immunofluorescence in whole embryos

Embryos were fixed using formaldehyde as describe in the section ‘Collection and fixation of embryos’. The methanol-stored embryos were rehydrated and washed in 1x PBS with 0.1% Tween-20 for 10 min. The immunofluorescence protocol was performed as previously described^55^ for anti-RBP1 conjugated with Alexa Flour 488 (CTD4H8, Sigma-Aldrich, 1:100 dilution), targeting all forms of RNA Pol II with the following modifications. Anti-RBP1 conjugated with Alexa Flour 488 was incubated overnight in 1x PBS with 0.1% Tween-20 at 4 °C. The embryos were washed once for 30 min in 1x PBS with 0.1% Tween-20, followed by Hoechst 33342 (1:1000, Invitrogen) staining. The embryos were mounted in SlowFade Gold antifade reagent (Invitrogen) for imaging.

### Generation of haploid embryos

Homozygous *maternal haploid* (*mh)* virgin females (obtained from y[1] w[a] mh[1]/FM7a, Bloomington stock number 7130) were crossed with wild-type males^75-77,79-81^. The F1 haploid embryos were collected and fixed for FISH/Immunofluorescence to capture the major wave of ZGA (nuclear cycles 14 and 15) in haploid embryos. The control diploid embryos were from the wild-type Oregon-R (OR).

### Transcription inhibition using microinjection

The transcription inhibition using microinjections was performed with modifications^6^. Virgin females from DGRP-057 and males from DGRP-439 lines were crossed and allowed to lay embryos for 45 min at 25 °C. Chorion was manually removed using double-sided sticky tape and forceps. Approximately 40-50 embryos were lined up on a clean microscope slide and covered with 50-100 µl of halocarbon oil 0.8 (Apollo Scientific). Based on the injection pressure on the Eppendorf FemtoJet 4i microinjector, approximately 0.2 nL of water (control), alpha-amanitin (0.5 mg/ml in water, Sigma-Aldrich) or triptolide (1 mg/ml in DMSO, followed by dilution in water to obtain 0.05 mg/ml, Selleck Chemicals) was injected into the embryos. Post injection, the embryos were incubated at 25 °C on the microscope slide. At 2.5 hours after egg laying (AEL), the embryos were gently removed from the slide using a brush, transferred into the formaldehyde-fix to continue with the fixation as described in the section ‘Collection and fixation of embryos’.

## Acknowledgments

We thank all members of the Erceg laboratory for discussion, Barbara G. Mellone for Oregon-R fly line, the Advanced Light Microscopy Facility at the University of Connecticut, and the Bloomington *Drosophila* Stock Center for *Drosophila* lines. We apologize to the authors whose work we could not include due to space constraints. T.M.O., R.R., and P.I.F. were supported by a Summer Undergraduate Research Fund (SURF) awards. Work in J.E.’s laboratory was supported by the University of Connecticut and an award to J.E. from NIH/NIGMS (R35GM146922). M.I.’s laboratory was supported by NIH/NIGMS (R35GM128678).

## Author Contributions

A.S.G. and J.E. designed research. A.S.G., T.M.O., R.R., P.I.F., A.J., and S.M.R. performed experiments with input in experimental design from M.I. and J.E. A.S.G. and A.Z. analyzed data. A.S.G. and J.E. wrote the manuscript. All authors approved the final version of this manuscript.

## Supplemental information

**Figure S1.**
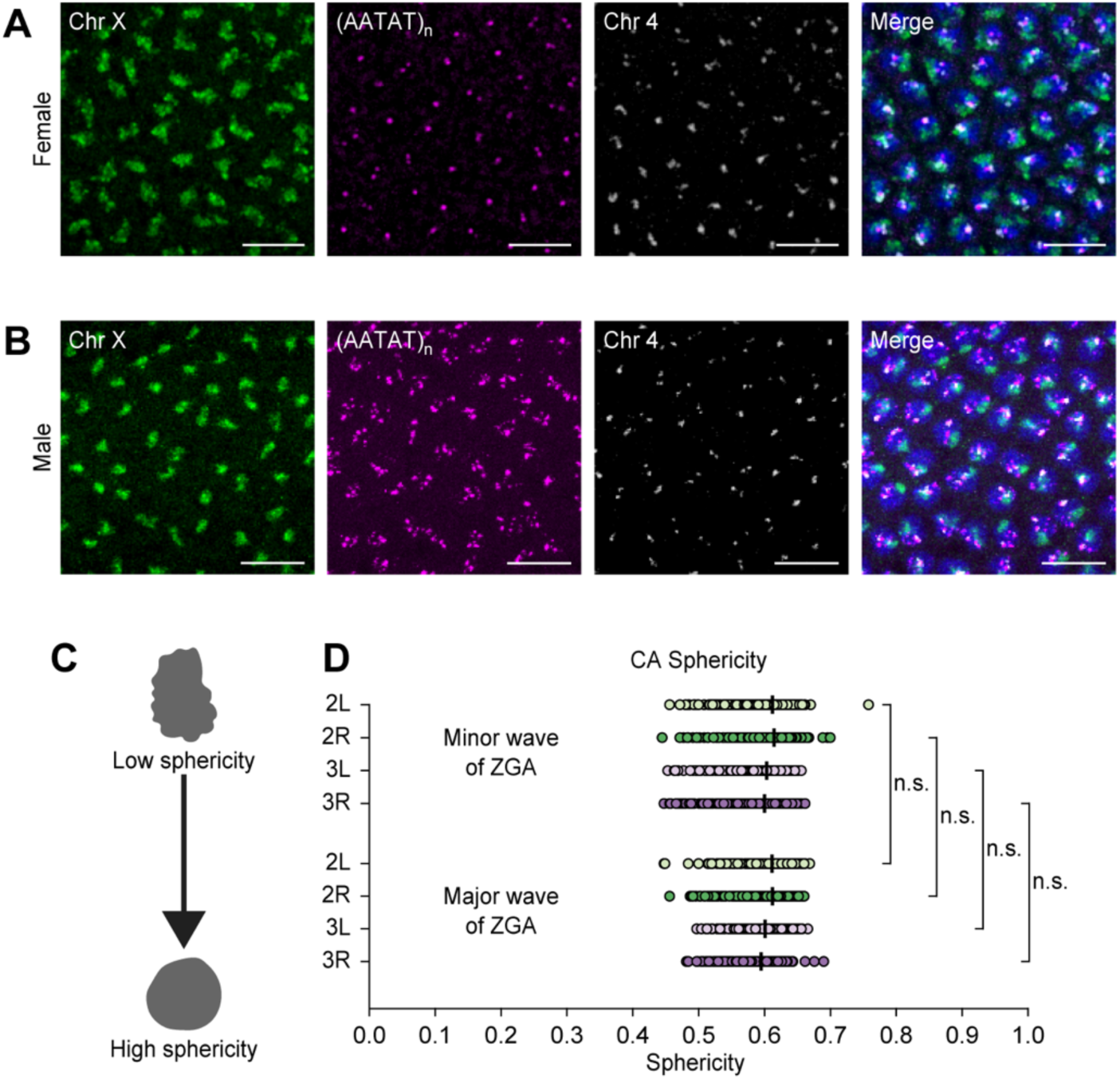
Chromosome X in female and male embryos and sphericity of CAs. (A and B) Chromosome X (green), satellite repeat probe (AATAT)_n_ (magenta), and chromosome 4 (gray) in female (A) and male (B) embryos during major wave of ZGA. Total DNA by Hoechst stain (blue). Bar = 10 μm. (C) Illustration showing low and high sphericity. (D) CA sphericity between the minor and major waves of ZGA. Median, solid line; at least three replicates; n ≥ 300 nuclei; n.s., not significant, Mann-Whitney two-sided *U* test.

**Figure S2.**
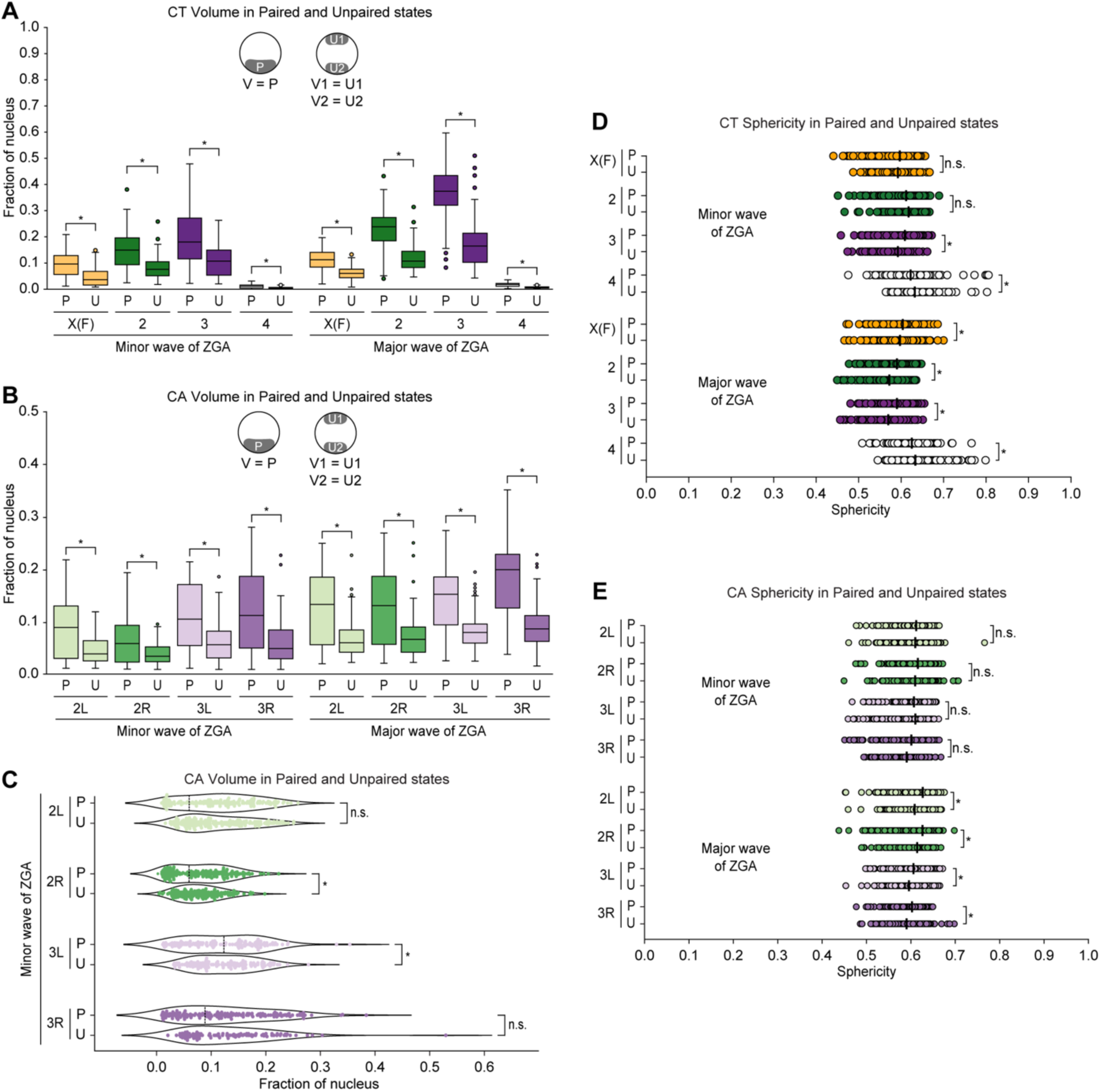
Volume and sphericity changes of CTs and CAs based on pairing. (A) Normalized CT volume changes between paired homologs to individual unpaired homologs from the minor to major waves of ZGA. X(F), chromosome X in females; P, paired; U, unpaired; U1, unpaired homolog 1; U2, unpaired homolog 2; V, volume; at least three replicates; n ≥ 128 nuclei; *p ≤ 1.26×10^-9^, Mann-Whitney two-sided *U* test. (B) Normalized volume differences in arms of paired homologs to arms of individual unpaired homologs during ZGA. P, paired; U, unpaired; U1, unpaired homolog 1; U2, unpaired homolog 2; V, volume; at least three replicates; n ≥ 109 nuclei; *p ≤ 2.67×10^-4^, Mann-Whitney two-sided *U* test. (C) Normalized CA volume differences in arms of paired homologs to the combined volume of two unpaired homologs during ZGA. P, paired; U, unpaired; local minima, dashed line; at least three replicates; n ≥ 103 nuclei; *p ≤ 1.12×10^-3^, n.s., not significant, Levene’s test. (D) CT sphericity changes between paired and unpaired homologs during ZGA. X(F), chromosome X in females; P, paired; U, unpaired; median, solid line; at least three replicates; n ≥ 148 nuclei; *p ≤ 3.58×10^-2^, n.s., not significant, Mann-Whitney two-sided *U* test. (E) CA sphericity of paired and unpaired homologs during ZGA. P, paired; U, unpaired; median, solid line; at least three replicates; n ≥ 150 nuclei; *p ≤ 3.77×10^-4^, n.s., not significant, Mann-Whitney two-sided *U* test.

**Figure S3.**
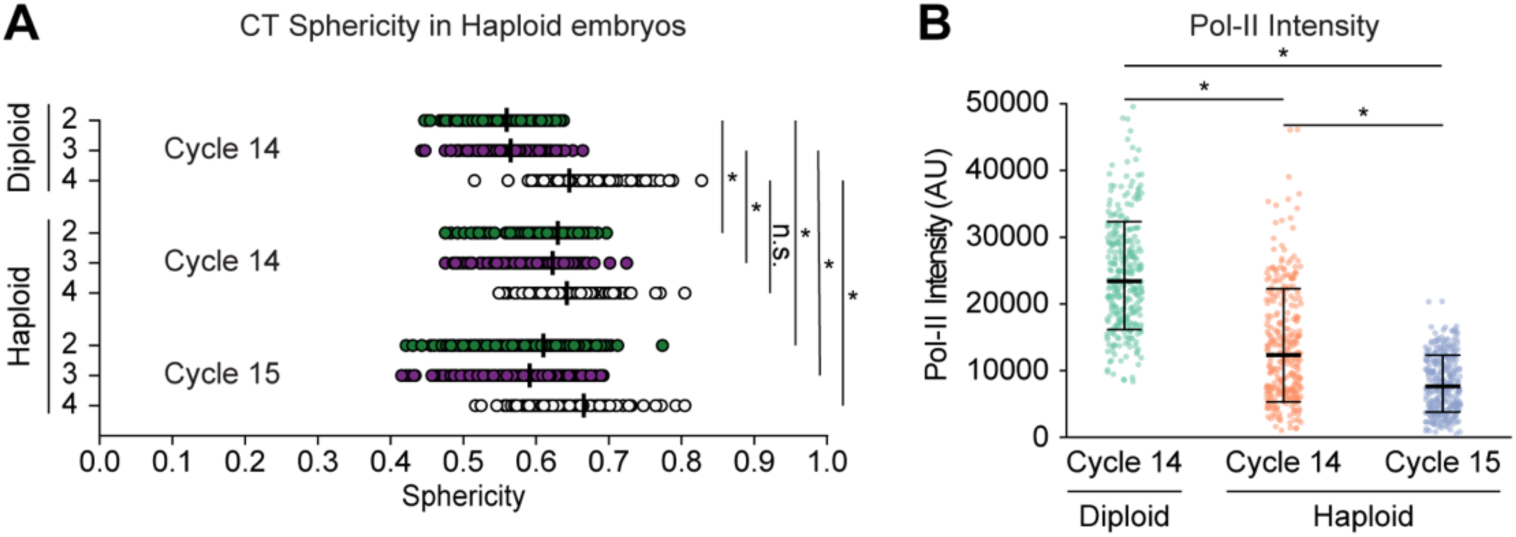
Sphericity of CTs and RNA Pol II intensity for diploid and haploid embryos. (A) CT sphericity in diploid and haploid embryos during major wave of ZGA. In diploid embryos, only individual unpaired homolog sphericity was used. Median, solid line; at least three replicates; n ≥ 300 nuclei; *p ≤ 1.87×10^-7^, n.s., not significant, Mann-Whitney two-sided *U* test. (B) Distribution of RNA Pol II fluorescence intensity (a.u.) within nucleus for diploid and haploid embryos during major wave of ZGA. At least three replicates; n ≥ 300 nuclei; *p ≤ 3.33×10^-18^, Mann-Whitney two-sided *U* test.

**Table S1.** Oligopaint summary and CT/CA changes during minor and major waves of ZGA. *See separate excel sheet*.

(A) Summary of Oligopaint probes designed. (B) Primers for Oligopaint probes. (C) Normalized CT volume and (D) sphericity during ZGA. (E) Normalized CA volume and (F) sphericity during ZGA. X(M), chromosome X in males; X(F), chromosome X in females.

**Table S2.** Pairing of CTs and CAs during ZGA. *See separate excel sheet*.

(A) CT pairing and (B) CA pairing during minor and major waves of ZGA. (C) Normalized CT volume and (D) CA volume changes between paired homologs to the combined volume of two unpaired homologs. (E) Normalized CT volume and (F) CA volume differences between paired and individual unpaired homologs. (G) CT sphericity and (H) CA sphericity between paired and unpaired homologs. P, paired; U, unpaired.

**Table S3.** CT and RNA Pol II dynamics in haploid embryos. *See separate excel sheet*.

(A) Nuclear volume, (B) normalized CT volume, and (C) HLB volume in diploid and haploid embryos during major wave of ZGA. (B and C) In diploid embryos, only individual unpaired homolog volumes were used. (D) CT sphericity differences in diploid and haploid embryos. In diploid embryos, only individual unpaired homolog sphericity was used. (E) RNA Pol II fluorescence intensity (a.u.) within nucleus for diploid and haploid embryos.

**Table S4.** CT and nuclear volume measurements in transcription inhibited embryos. *See separate excel sheet*.

(A) Nuclear volume, (B) normalized CT volume, and (C) CT pairing in transcription inhibited embryos during major wave of ZGA.

